# “Spatial pattern of herbaceous seed dispersal by ungulates in grasslands of the Doñana National Park (SW Spain)”

**DOI:** 10.1101/2025.06.19.660649

**Authors:** María José Leiva, José María Fedriani

**Author notes:** **Corresponding author -** María José Leiva.

## Abstract

Most research on endozoochorous seed dispersal by ungulates emphasizes long-distance dispersal and its dependence on ungulate species and habitat heterogeneity. In contrast, the role of ungulate community composition in shaping short-distance dispersal and generating spatial patterns in the distribution of plant diaspores remains largely unexplored— key questions addressed in this study. We conducted fieldwork at two adjacent grassland sites in Doñana National Park (southwestern Spain). One site was used by a mixed community of four ungulate species—both wild and domestic (deer, wild boar, cattle, and horses)—while the other was used almost exclusively by deer. Our results revealed clear differences between sites in the total number of dispersed herbaceous seeds and the most frequently dispersed plant families. Within the four-species community, cattle and deer differed most in the taxonomic composition of the seeds they dispersed, suggesting that herbivore-specific seed selection and dispersal act as key drivers of grassland structure at fine spatial scales, mirroring dynamics typically observed at broader scales. Moreover, only in the four-ungulate community did we detect significant spatial patterns in seed dispersal. These included positive effects of short-distance feces aggregation, spatial covariance in seed content among nearby feces, and a strong correlation between seed content and local feces density. Seed families predominantly dispersed by cattle also exhibited significant spatial structuring. These findings have important implications for biodiversity management in semi-natural and protected ecosystems, underscoring the need to consider functional differences among herbivores when developing conservation and restoration strategies.

## 1. INTRODUCTION

Large herbivorous ungulates are important seed dispersers through both endozoochory and epizoochory. Thus, these dispersers contribute to the formation of plant seed banks, particularly for herbaceous species (Boulanger et al. 2011; Pellering et al. 2016), while also playing a key role in shaping herbaceous plant communities (Janzen 1984; Milchunas et al. 1988; Nathan and Muller-Landau 2000; Cosyns et al. 2005). The guild of ungulate species coexisting in an area is frequently considered functionally redundant in terms of seed dispersal (O’Connor and Crowe 2005), because most large ungulates consume a wide variety of herbaceous plants (Janzen 1984; Baltzinger et al. 2019; Delibes et al. 2019). However, low redundancy in seed dispersal has also been observed among sympatric ungulates. For instance, Picard et al. (2016) found that in agro-forested landscapes, the feces of red deer (Cervus elaphus) contained greater seed abundance and species richness than those of sympatric wild boar (Sus scrofa) and roe deer (Capreolus capreolus). Polak et al. (2014) also identified a lack of redundancy in seed dispersal by two reintroduced wild ungulates — the Arabian oryx (Oryx leucoryx) and the Asiatic wild ass (Equus hemionus) — in the Negev Desert, where another ungulate, the dorcas gazelle (Gazella dorcas), has historically been present. In this case, the three species of ungulates dispersed different assemblages of plant species.

Ungulates are key long-distance seed dispersers (Albert et al. 2015; Pellerin et al. 2016; Nathan and Muller-Landau 2000). Thus, their role in maintaining habitat connectivity through seed dispersal in fragmented landscapes (Cosyns et al. 2005; Moore and Swihart 2007; Pakeman and Small 2009; Pellerin et al. 2016) and the differences in habitat selection and plant dispersal among ungulate species have frequently been documented, particularly in large areas composed of a mosaic of land uses and vegetation types (Albert et al. 2015; Picard et al. 2016). Studies of seed dispersal by ungulates at medium and small spatial scales have also been developed in recent decades, focusing mainly on the degree of convergence between local flora and dispersed seeds (Cosyns et al. 2005), temporal changes in seed dispersal within local herbaceous communities (Malo 1995b), or differences among local ungulates in the pool of dispersed seeds (Bartuszevige and Endres 2008), among others. However, studies on the spatial patterns of seed dispersal mediated by ungulate herbivores in grassland communities remain much scarcer (but see Malo et al. 2000). Importantly, spatial structure is critical for determining the dynamics of plant communities (Coughenor 1991), and short-distance seed dispersal plays a key role in shaping the potential area of plant recruitment and subsequent ecological processes such as competition, mating, and predation (Nathan and Muller-Landau 2000).

In the last century, wild ungulate populations have experienced significant density increases in many areas of Europe due to the reduction of natural predators and changes in management practices (e.g. supplementary feeding) (Acevedo et al. 2011, Lecomte et al. 2016). Additionally, wild ungulates often coexist with domestic ungulates in many human-managed grazing areas (Malo and Suarez 1995b, Kukielka et al. 2013, Tapia-Excárate et al. 2021, Hines et al. 2021), including protected areas where domestic species may still persist as remnants of past land use (Garrote et al. 2022, OAPN 2025). However, the effects of ungulate community composition on seed dispersal and spatial pattern generation at the local grassland scale remain largely understudied.

In this study, we assess the dispersal of herbaceous seeds by two ungulate communities within protected Mediterranean grasslands in Doñana National Park (southern Spain). This area is recognized as one of Europe’s most important strongholds for biodiversity conservation (Martín-López et al. 2007) and has been the focus of extensive research on seed dispersal mediated by birds and frugivorous mammals (Coughlan et al. 2017, Perea et al. 2013, Fedriani et al. 2010, Green et al. 2022, Fedriani et al. 2023), frequently at the scale of single plant populations (Fedriani et al. 2024). However, very few studies have addressed seed dispersal by ungulate herbivores in the park’s grassland ecosystems, where numerous herbaceous species may be dispersed endozoochorously under the so-called “foliage is the fruit” hypothesis (Janzen 1984). The objectives of this study are i) to assess potential differences in the quantity and quality (taxonomic composition) of herbaceous seeds endozoochorously dispersed by ungulates in two nearby grassland sites that differ in ungulate species richness and composition, as well as in the domestic or wild status of these ungulate communities, and ii) to investigate potential spatial patterns of herbaceous seed dispersal generated by these two communities, both in terms of total seed number and the most frequently dispersed seeds taxa. Based on findings from other European habitats (Jaroszewicz et al. 2013), we hypothesize that the two ungulate communities will differ in both the amount and the taxonomic composition of dispersed seeds. Furthermore, we expect differences in the spatial pattern of seed deposition, reflecting variation in ungulate size and behavior (Coughenour 1991, Malo et al. 2000).

## 2. MATERIAL AND METHODS

### 2.1 Study area and sites description

The study was conducted in Doñana National Park, located in southern Spain (510 km²; 37° 10′ N; 6° 26′ W) within the Guadalquivir River estuary — a flat area ranging from 0 to 80 m above sea level (Fedriani & Wiegand 2014). The climate is Mediterranean, with wet, mild winters and long, dry summers. The mean annual temperature is 17.7°C, while annual rainfall averages 465.6 ± 34.5 mm, although it varies considerably between years. Rainfall predominantly occurs from September to May and is scarce from June to August (data from ICTS Doñana, Palacio Meteorological Station, 2010–2022). The Doñana landscape is humanized and fragmented, with patches of woody vegetation frequently isolated by croplands, grasslands, marshes, dunes, and urban areas. The marshland, which is inundated during part of the year, is internally heterogeneous due to variation in elevation, flooding, and physical-chemical conditions (Soriguer-Escofet et al. 2001). Scrubland, another main habitat, typically occurs adjacent to the marshland in slightly higher, rarely flooded areas. It is also heterogeneous and supports a rich diversity of habitats (Fedriani & Wiegand 2014; Jiménez & Díaz-Delgado 2015).

In the contact zone between scrubland and marshland lies a transitional area predominantly composed of grassland, where this study was conducted. This ecotone, locally called “la vera”, varies considerably in width — from a few meters to several hundred — and is intensively grazed by ungulates (Villarfuertes et al. 1997; Barasona et al. 2014a, b). The ecotone is associated with a gradient of environmental conditions (such as water-table depth, soil texture, salinity, and pH; Lazo Contreras et al. 2001) that drives variation in the herbaceous community. Two main grassland types can be distinguished: dry grassland and wet grassland. The former occurs on sandy soils and is typically free from winter flooding. Representative species include the annuals *Anthoxanthum ovatum* and *Vulpia membranacea*, the perennials *Cynodon dactylon* and Panicum repens, and isolated individuals of *Asphodelus ramosus* and *Armeria gaditana*. Wet grassland develops on clayey-sandy soils, barely raised above the marsh level, and is prone to episodic winter flooding. The perennial *C. dactylon* is also abundant here and frequently occurs alongside dense stands of the reed *Juncus maritimus*. This ecotone is functionally connected to adjacent habitats; ungulates use the grasslands, the herbaceous gaps within scrubland, and the elevated areas of the marsh at different times of the year (Lazo Contreras et al. 2001).

Ungulates commonly found in the grassland include red and fallow deer (*Cervus elaphus* and *Dama dama*), wild boar (*Sus scrofa*) and domestic cow (*Bos taurus*) and horse (*Equus ferus caballus*). Their feeding habits vary seasonally (Soriguer et al. 2001). Generally, ungulates predominantly consume herbaceous plants from late winter to early summer but shift their diet toward woody species from mid-summer to early winter, when grassland biomass is scarce (Garrote et al. 2023 and references herein; Bugalho & Milne 2003; Schoenbaum et al. 2017 for other Mediterranean areas).

To conduct this study, two grassland sites were selected within “la vera”—Martinazo and Matasgordas, respectively. The sites are located approximately 10 km apart and each comprises both dry and humid grassland areas. Native vegetation at both sites has been altered by a long history of intensive human use, including livestock ranching, cultivation, and timber exploitation (Garrote et al. 2023). In Matasgordas, a large pastureland with patches of Mediterranean scrub and scattered *Quercus suber* and *Olea europaea* var. *sylvestris* trees, was created in 1970 by mechanically removing most shrubs and trees. In 1996, this area was classified by the Spanish National Park Services as a strict nature reserve, and livestock were largely removed from the site. Martinazo has a more extensive and prolonged history of intensive land use. Activities such as extensive livestock farming (cattle, horses, and pigs), wood gathering, and occasional cultivation following the felling of most trees have been practiced there since at least the 14th century (Fernández-Alés & Muñoz Reinoso 2020). The area was designated a Biological Reserve in 1964, which resulted in a partial reduction of human use, including a lower density of livestock. Today, Martinazo consists of a moderately extensive pastureland with a few sparse *Q. suber* and *O. europaea* var. *sylvestris* trees and scattered scrub patches. Livestock ranching (cattle and horses) still occurs there, although its intensity has diminished considerably in recent decades (Garrote et al. 2022). Currently, deer are 2.2 times more abundant in Matasgordas than in Martinazo, while cattle and horses are present only in Martinazo and are rare or absent in Matasgordas (Garrote et al. 2023). The two sites also differ in their physical characteristics. The topsoil at Martinazo contains approximately twice the amount of organic matter, phosphorus, and nitrogen compared to Matasgordas. Additionally, the groundwater table is shallower at Martinazo (2.39 m) than at Matasgordas (4.56 m) (Garrote et al., 2022).

At each site, a tetragonal plot was established for the collection of ungulate feces. The plots measured 10 ha at Martinazo and 90 ha at Matasgordas and were placed to minimize proximity to adjacent scrubland and marshland.

### 2.2 Feces sampling

Herbivore feces were sampled from early spring to mid-summer of 2020, corresponding to the fruiting season of most herbaceous species in the area (Valdés Castrillón et al., 1987). Sampling followed the methodology described by Fedriani et al. (2010). A series of starting points were regularly distributed along one edge of each plot. From each starting point, a non-systematic zigzag transect was conducted toward a non-fixed point on the opposite edge of the plot, and the return path followed a different trajectory. During each survey (i.e., a complete transect across the plot and back), all intercepted ungulate fecal units—defined as any spatial aggregation of fecal material such as pellet groups, dung pads, etc.—were recorded (deposition points hereafter). The spatial coordinates of each point were logged using a Global Position System-reading. Feces were identified to the species level based on morphological characteristics. Deer feces (including both red and fallow deer) are 1– 2.5 cm long and 0.8–1.4 cm wide, brownish-black, and of firm consistency; due to considerable overlap in size and shape, feces from both species were not reliably distinguishable and are hereafter referred to collectively as “deer.” Wild boar feces consist of compact units approximately 5 cm in diameter, often forming larger masses, dark brown to black in color, and with heterogeneous content. Horse feces are oval, at least 4 cm thick, smaller than a tennis ball, greenish-brown, and oblong with rounded ends, occurring singly or in amorphous groupings. Cow feces are larger, of pasty consistency, and typically deposited in flat, circular piles (Bang & Dahlström, 1992; Purroy & Varela, 2003).

In addition to feces recording, the presence or absence of woody vegetation near each deposition point was assessed to characterize the vegetation physiognomy of seed arrival microsites (e.g., García-Cervigón et al., 2017). A circular area with a 1-meter radius centered on each fecal unit was established, and the relative shrub cover was visually estimated using a subjective scale ranging from 5% to 100%.

Samples were collected from each fecal unit (50 cm³ per sample, when possible) and stored in labeled paper envelopes indicating the herbivore species, site, transect code, and date. Samples were kept at room temperature in the laboratory of the University of Seville until seed extraction. A total of 231 fecal samples were collected—113 from Martinazo and 118 from Matasgordas—during 145 surveys (65 in Martinazo and 75 in Matasgordas) conducted over 39 field visits. One sample (designated S72) contained 2,614 seeds, representing 37% of the total recorded seeds. Due to its disproportionate contribution, this sample was excluded from all analyses to avoid skewing the results.

### 2.3 Seed identification and quantification

Approximately 2.5 g of dry weight (DW) per sample were used for seed extraction, except in cases with insufficient material (16% of samples, *n* = 37), where the entire sample was processed. The dry weight of each individual sample was recorded prior to processing. Samples were rehydrated and carefully washed using a series of sieves with mesh sizes of 0.5 mm, 0.1 mm, and 0.01 mm under running water. The retained material was then visually inspected under a stereo microscope (8×–35×, Leica EZ4) to detect the presence of seeds. Only unbroken, non-empty, and apparently intact seeds were selected and stored in Petri dishes for subsequent identification. The dry weight of each sample was used to standardize seed content, expressed as the number of seeds per gram DW.

Seeds were identified to the highest possible taxonomic level using *Flora Ibérica* (Castroviejo, 1986–2012) and *Flora de Andalucía Occidental* (Valdés Castrillón et al., 1987). Additionally, a reference seed collection, compiled during the study year from plant species occurring at the study sites, was consulted. Expert taxonomists also collaborated in identifying specific seed taxa, including Dr. Benito Valdés Castrillón and Dr. Ádám Lovas-Kiss.

In addition to standardizing seed content by weight, seed content was also estimated on a per fecal unit basis. This approach was necessary given the substantial variation in fecal mass among herbivore species (deer, cow, horse, and wild boar), each of which produces different quantities of dung per defecation (Appendix S1, Table 1, and references therein). Thus, herbivore species likely represent a major source of variation in seed deposition via feces, a factor particularly relevant to understanding spatial seed dispersal patterns (Perea et al., 2012; Zwolak, 2018). To account for these differences, the number of seeds potentially deposited at each deposition point was calculated as the product of the standardized seed content (seeds per gram DW) and the average dry weight per defecation for each species, based on values from published sources (Appendix S1, Table 1).

### 2.4 Data analyses

#### 2.4.1 Taxonomic composition of dispersed seeds. Effect of site and of ungulate species

To evaluate the potential effect of site (i.e., Martinazo vs. Matasgordas) on the taxonomic composition of dispersed seeds, a non-metric multidimensional scaling (NMDS) analysis was performed using Euclidean distance as the similarity index. Seed content was expressed both per gram DW and per fecal unit. Additionally, a PERMANOVA (Permutational Multivariate Analysis of Variance) was conducted to detect the significance of differences. This analysis included only data from the ungulate species common to both sites (i.e., deer), excluding the two wild boar samples and the outlier sample S72 from Matasgordas, and including only deer samples from Martinazo (representing 48.7% of the total samples from that site). NMDS based on Euclidean distance was also applied to assess potential differences in the taxonomic composition of dispersed seeds among the four ungulate species. PERMANOVA was again used to test for overall significant differences among species, and Bonferroni correction was applied for pairwise comparisons between them. All statistical analyses were conducted using the PAST v4.17 software package.

#### 2.4.2. Spatial pattern of seed dispersal

Techniques of marked point pattern analysis (Illian et al. 2008; Wiegand and Moloney 2014) were used to evaluate potential spatial patterns in herbivore-generated seed rain at Martinazo and Matasgordas. The approach was applied first to the total number of dispersed seeds and then to those from the most frequently dispersed plant families that were present in a sufficient number of samples to allow for robust analysis. The datasets comprised spatial coordinates (*xi*) and marks (*mi*) representing seed content (i.e., number of seeds). Seed content is expressed per fecal unit to provide a more realistic representation of seed arrival in the presence of different ungulate species. The data structure consists of a univariate quantitatively marked point pattern, where the coordinates (*xi*) represent the univariate point pattern and the seeds content (*mi*) represent a quantitative attribute (i.e., mark).

Mark correlation functions are based on all (ordered) pairs of fecal units whose inter-point distances lie within a small interval (*r* − *h*, *r*+ *h*). The parameter *h,* called the bandwidth, must be sufficiently large to provide an adequate number of pairs in each distance class “*r”* but small enough to resolve relevant biological details (Illian et al., 2008). The objective of mark correlation functions is to estimate the mean value *c*_t_(*r*) of a test function *t*(*m_i_*, *m_j_*) for two marks *m_i_* and *m_j_*, taken over all (ordered) pairs of fecal units *i* – *j* with inter-point distances of *r*± *h*.

This procedure is then repeated across a range of distances *r,* using intervals Δ*r* to produce the non-normalised mark correlation function *c*t(*r*) (Illian et al., 2008). To obtain the final mark correlation function, *c*t(*r*) is normalized by the expected value *c*t of the test function taken over all pairs of fecal units, regardless of their spatial separation:

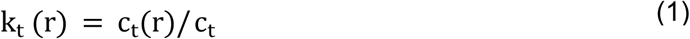

Many different test functions *t*(*m_i_*, *m_j_*) are possible. Here we use three of them which are complementary in the information they provide (Fedriani et al. 2015; Wiegand and Moloney 2014). Specifically, we used the *r*-mark correlation function *k*_m_(*r*), which is based on the test function:

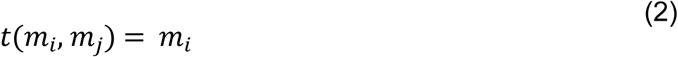

The estimator of the corresponding non-normalised mark correlation function is given by:

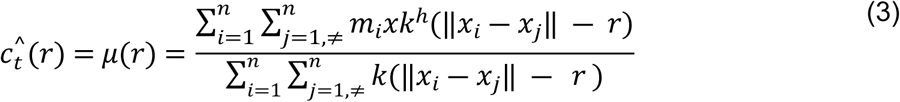

Where the “box kernel” function *k^h^(d)* yields a value of 1/2*h* if the two fecal units with coordinates x*i* and x*j* are at a distances of *r*± *h*, and zero otherwise (Illian et al., 2008; Wiegand & Moloney, 2014). Thus, the denominator of Eq. (3) corresponds to the number of ordered pairs of fecal units *i* and *j* that are distances *r*± *h* apart; Eq. (3) calculates the mean mark *m_i_* of the first fecal units *i* of these pairs. The normalisation constant *c*_t_ of the *r*-mark correlation function, which is taken over all pairs of points regardless of their spatial separation, is obtained by replacing *k^h^*(*d*) in Eq. (3) with 1/2*h*:

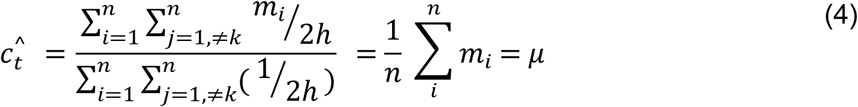

and yield μ, the mean value of m_i_ taken over all fecal units *i* (Illian et al. 2008).

Thus, for the r-mark correlation function, *km(r)>1* indicates a positive effect of feces aggregation (i.e., the seed content of feces that have nearby feces at distance *r* is on average larger than the average seed content across feces), while *km*(*r*) < 1 indicates a negative effect (i.e., the seed content of feces that have nearby feces at distance *r* is on average smaller than the average seed content) (e.g. Fedriani et al., 2015).

We also used a mark correlation function that characterizes the spatial covariance in seed number of two fecal units separated by distance *r*. This test function is known as Schater’s and was proposed by Schlather, Ribeiro and Diggle (2004):

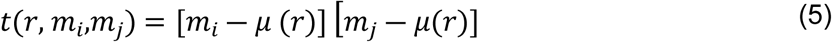

It results in a Moran’s I-like summary statistic I_mm_ (r), a spatial variant of the classical Pearson correlation coefficient (Shimatani, 2002). I_mm_ (r) is normalized by mark variance σ^2^. Thus, I_mm_ (r) is the unequivocal Pearson correlation coefficient between the two variables m_i_ and m_j_ defined by the ordered i-j pairs of feces separated by distance *r*± h. Note that a test function that adjusts for the mean μ(*r*) that considers only pairs of fecal units separated by a given distance *r*, not the population mean μ, is required to get a summary statistic that can be interpreted as a correlation coefficient (Schlather et al., 2004).

Finally, we use the density correlation function *Cm,K*(*r*) developed by Fedriani et al. (2015), which directly relates the seed content in a fecal unit to the density of its nearby feces. *C_m,K_*(*r*) estimates the classical Pearson correlation between the seed content *m_i_* of a fecal unit and the number of nearby feces within distance *r* [=λ*K_i_*(*r*)]. Thus, the density correlation function is based on the following test function:

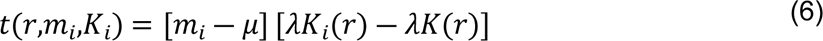

where *m_i_*is the seed content of focal fecal unit *i*, μ is the mean seed content of the feces population, λ is the overall density of feces in the study area, λ*K_i_*(*r*) is the number of nearby feces around focal fecal unit *i* within distance *r* and λ*K*(*r*) is the mean number of nearby feces within distance *r* for all fecal units.

## 3. RESULTS

### 3.1 Vegetation physiognomy at the seed arrival microsites, and importance of different dispersers at each site

As expected, vegetation physiognomy at the seed arrival microsites in both Martinazo and Matasgordas was predominantly composed of open grasslands (78-82% microsites were exclusively covered by grassland; Table 1) while 18 to 22% microsites were partially occupied by shrubs (= shrubby microsites) with moderate (28-31%) cover. *Halimium halimifolium* was the dominant shrub species in most shrubby microsites at both locations, although the second most frequently dominant species differed: *Stauracanthus genistoides* in Matasgordas and *Asparagus acutus* in Martinazo.

**Table 1.**
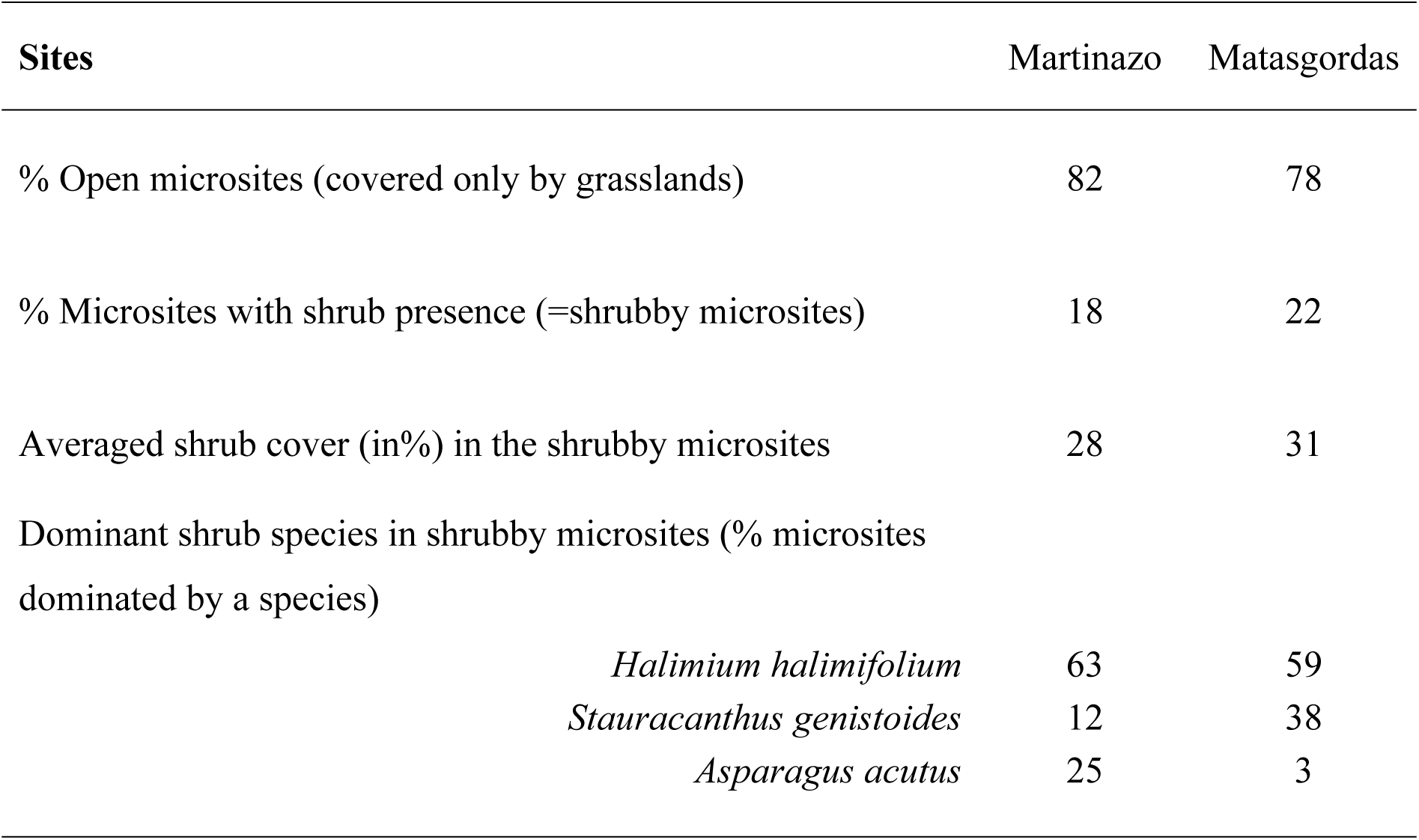
Vegetation physiognomy in seed arrival microsites.

As anticipated, only wild ungulates feces were found in Matasgordas (Table 2), with deer strongly dominant (98.3% feces) and wild board negligible (2 feces; 1.7% of total). In Martinazo, deer were also most abundant (49% feces), followed by cattle (27 % feces) and horses (12% feces) and wild board (12% feces) in much lower proportion. Overall, fecal density was considerably lower in Matasgordas (1.3 feces.ha^−1^) than in Martinazo (11.4 feces.ha^−1^).

**Table 2.**
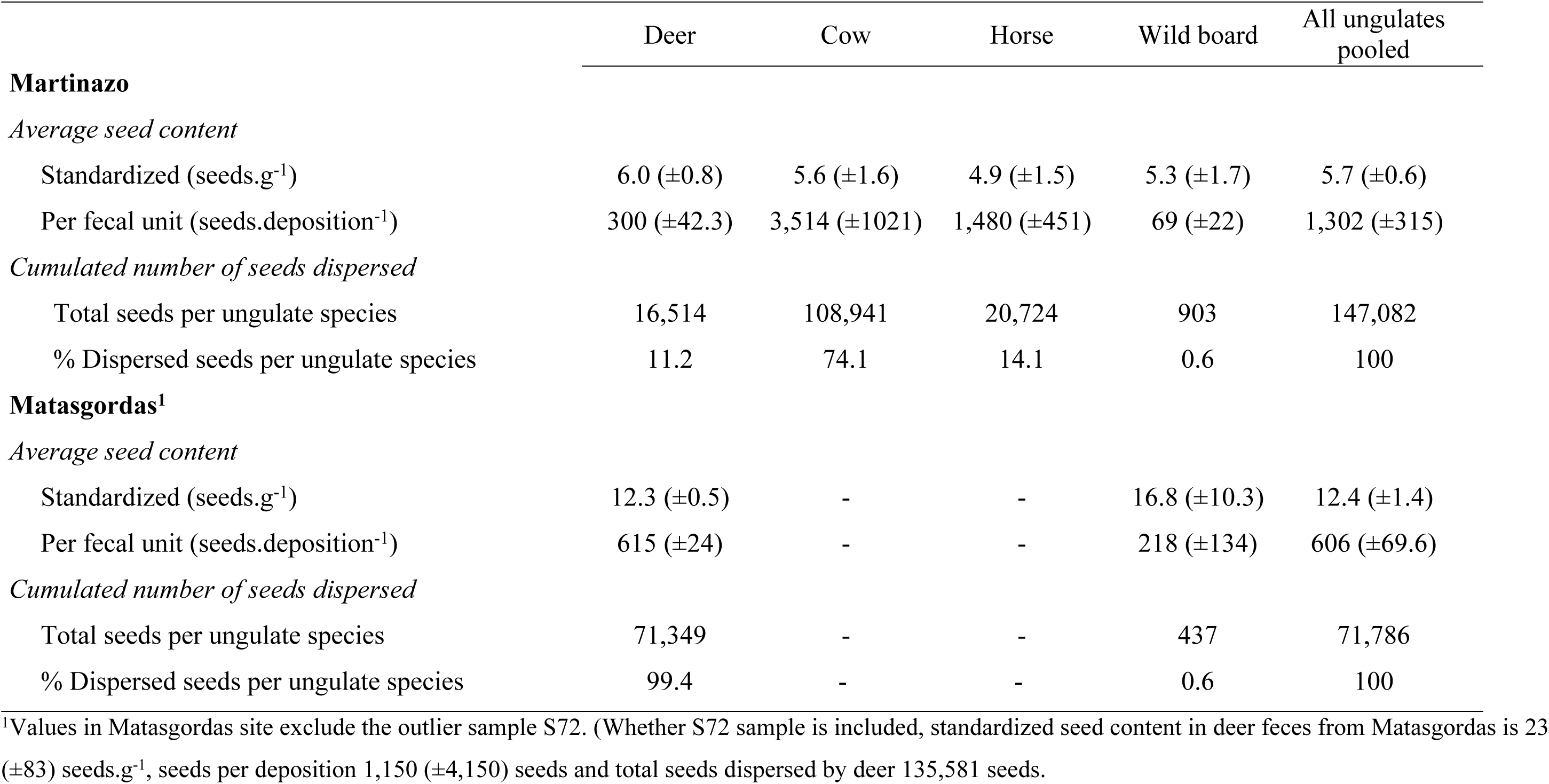
Seed content (expressed per gram dry weight and per fecal unit DW) in depositions from different ungulates at Martinazo and Matasgordas (mean values ± se). The seed content per fecal unit is calculated from average feces DW of different ungulates (Appendix S1 Table1). Cumulated number of dispersed seeds is the sum of seeds in all depositions from each ungulate.

Overall seed content varied from 0 to 97 seeds.g^−1^ across all samples, excluding the outlier S72 (Table 2). Seed content per gram DW was generally higher in Matasgordas than in Martinazo (Table 2), but the opposite pattern was observed when expressed per fecal unit. This reflects the nearly exclusive presence of deer in Matasgordas which produce lighter droppings with lower seed content than cattle and horses (Appendix S1 Table1), species that, alongside deer and wild boar, are present in Martinazo.

### 3.2 Taxonomic composition of dispersed seeds. Effect of site and ungulate species

Seeds extracted from fecal samples were predominantly small (<1mm), rounded, and or hard-coated. Of the total 7,074 seeds recovered, only 233 (3.3%) could not be identified, while the rest were identified to various taxonomic levels. In total, 21 families, 53 genera, and 57 species were identified (Appendix S1 Table 2). Family was determined for the majority (96.7%) of seeds and was therefore selected for subsequent analysis of taxonomic composition.

The importance (i.e., relative abundance and frequency of occurrence) of different plant families in the seeds extracted from Matasgordas and Martinazo samples (excluding S72) is shown in Figure 1. The large majority of dispersed plant families were present at both sites (i.e., 17 families, Figure 1a) while three were exclusive to Matasgordas (i.e., *Chenopodiaceae, Plumbaginaceae, Euphorbiaceae*) and one exclusive to Martinazo (*Alismateceae*). Exclusive families were scarce at their respective sites with abundances ranging from 0.49 to 0.07%. For the most abundant families, the abundance ranking varied between the two sites. The most abundant dispersed family in Matasgordas (*Valerianaceae,* 37% seed abundance, Figure 1a) was very scarce in Martinazo (1.9% *Valerianaceae* seed abundance). This family was represented by a single species (*Centranthus calcitrapae* Appendix S1 Table 2). Similarly, the second and third most abundant families in Matasgordas, (*Fabaceae,*16% abundance; *Juncaceae* 15%) were moderately abundant or scarce in Martinazo (*Fabaceae,* 8.2% abundance; *Juncaceae* 1.4 % seed abundance). The *Juncaceae* family was represented by a single genus, *Juncus* (Appendix S1 Table 2) which was also the only genus present in the outlier sample S72. Conversely, the three most abundant dispersed families in Martinazo (*Plantaginaceae,* 19%; *Cyperaceae,* 17%; *Ranunculae*, 14 % seed abundance) were scarce in Matasgordas (*Plantaginaceae*, 3%; *Cyperaceae,* 4%; *Ranunculae,*5% seed abundance). It is also noteworthy that seed abundance was fairly unevenly distributed across families in Matasgordas (Figure 1b), where the most abundant family *(Valerianaceae)* was considerably more abundant than the next two (*Fabaceae* and *Juncaceae).* In contrast, seed abundance in Martinazo was more evenly distributed (Figure 1c) with the three most abundant families (i. e., *Plantaginaceae, Cyperaceae* and *Ranunculaceae)* occurring in similar proportions.

**Figure 1.**
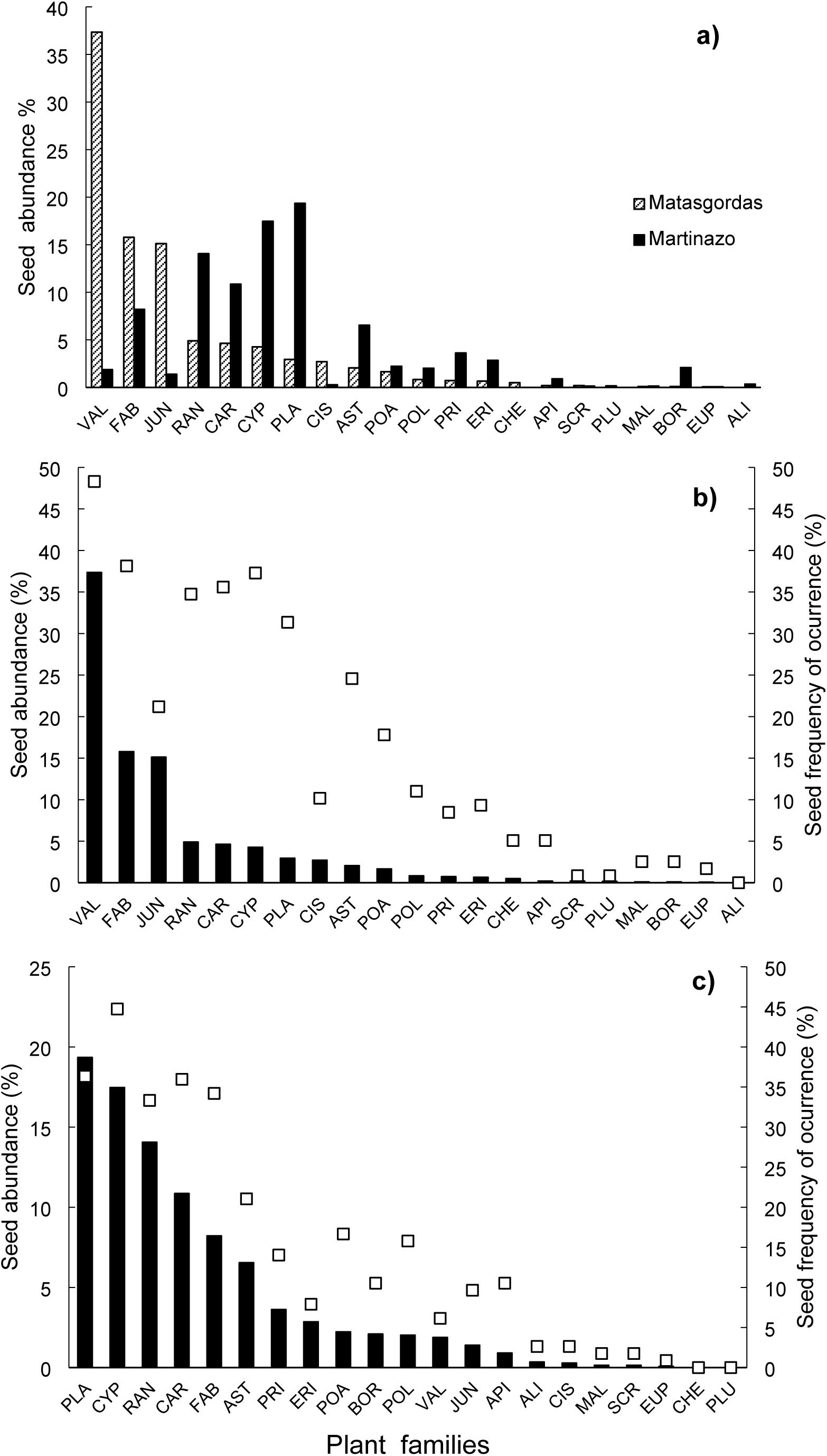
**(a)** Relative abundance of different plant families (percentage of the seed pool) at the estudied sites, and combined relative abundance of plan families (= black bars) and their frecuency (percentage of samples where a family occurs = white squares) in Matasgordas **(b)** and Martinazo **(c)** The sample S72 is excluded from computation. Plant families: VAL (*Valerianaceae*), FAB (*Fabaceae*), CYP (*Cyperaceae*), PLA (*Plantaginaceae*), RAN (*Ranunculaceae*), CAR (*Caryophyllaeae*), AST (*Asteraceae*), CIS (*Cistaceae*), POA (*Poaceae*), PRI (*Primulaceae*), ERI (*Ericaceae*), POL (*Polygonaceae*), BOR (*Borraginaceae*), API (*Apiaceae*), CHE (*Chenopodiaceae*), SCR (*Scrophulariaceae*), MAL (*Malvaceae*), ALI (*Alismataceae*), PLU (*Plumbaginaceae*), EUP (*Euphorbiaceae*).

The discrepancy between the frequency of occurrence and relative abundance of dispersed families was more was particularly pronounced at Matasgordas (Figure 1b), where several families exhibited disproportionately high frequency relative to their relative abundance (e.g., *Fabaceae, Ranunculaceae, Caryophyllaceae, Cyperaceae, Plantaginaceae, Asteraceae, Poaceae*). At Martinazo (Figure 1c), only a few families —namely *Fabaceae* and *Caryophyllaceae* showed marked discrepancies. Notably, *Fabaceae* and the *Caryophyllaceae* consistently exhibited high discrepancies at both sites, being frequently dispersed despite their low relative abundance. This suggests that many dispersers consume seeds from these families, albeit in small quantities.

NMDS results comparing the taxonomic composition of the dispersed seeds by deer at Matasgordas and Martinazo (Figure 2) show a high degree of overlap in family composition. However, several seed families were more prominetly represented in Matasgordas (e.g., *Valerianaceae, Juncaceae, Fabaceae, Chenopodiaceae and Cistaceae),* while others were better represented in Martinazo (e.g., *Ranunculaceae* and *Borraginaceae*). PERMANOVA analysis indicated signifficant differences between the two sites (Df =1, F = 7.01, *P* = 0.0001). In this anlysis, seed content was expresed per gram of dry weight. Nonetheless, when expressed per faecal unit, results remained consistent (data not shown) as above, as both data sets are proportional due to the involvement of the same disperser species (deer).

**Figure 2.**
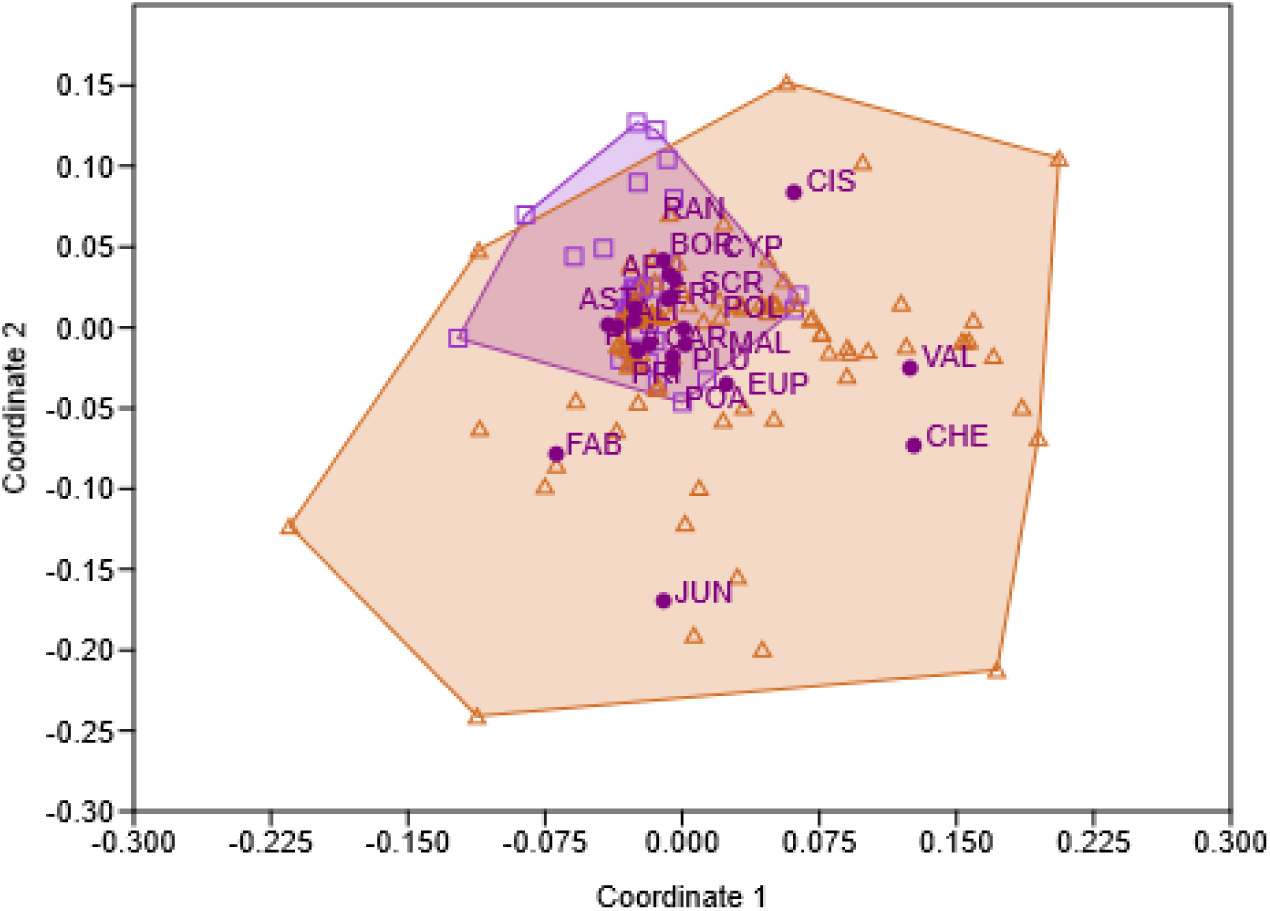
NMDS results showing deer samples from Martinazo and Matasgordas sites The analysis is based on seeds per gram DW. Squares and purple area = Martinazo deer, triangles and pink area = Matasgordas deer. Initials next to dots are seed families positioned by abundance and frequency in each site. Initials of family names as in Figure 1

NMDS comparing taxonomic composition of seeds dispersed by different ungulates at Martinazo are presented in Figure 3. As in the previous case, there was a high degree of overlap in seed composition when data were expresed per gram DW (Figure 3a). Nevertheless, certain plant families were preferenttially associated with specific dispersers. Deer and cattle were the most dissimilar ungulates in the taxonomic composition of dispersed seeds, with *Ranunculaceae*, *Polygonaceae and Borraginaceae* families more closely associated with deer, and *Plantaginaceae* with cattle. Wild board and horse occupied intermediate position: wild board showed strong overlap with deer and partial overlap with cow and horse, while horse overlapped closely with both deer and cow. PERMANOVA analysis (Table 3, column A) revealed significant overall differences among ungulate species (Df = 3, F = 2.142, P = 0.012). However, pairwise comparisons using the Bonferroni-corrected test indicated that only the deer–cow comparison was statistically significant.

**Figure 3.**
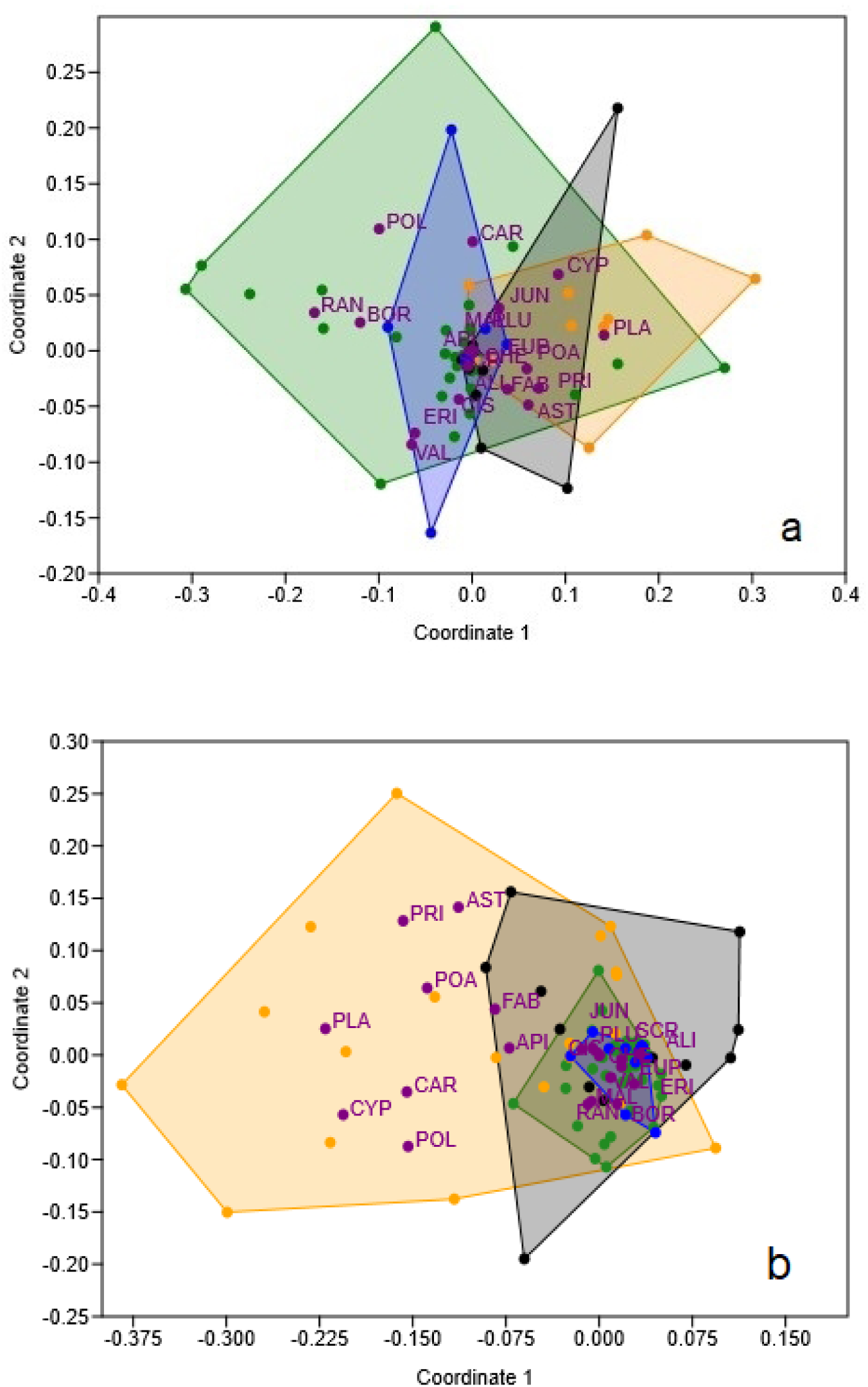
NMDS results showing samples of different ungulate species (dots and areas of different colours) at Martinazo: Green = deer; yellow = cow; black = horse, blue = wild boar. Seed content expresed per gram DW **(a)** and per fecal unit **(b).** Initials next to dots are families of seeds positioned by abundance and frequency in each ungulate. Initials of family names as in Figure 1.

**Table 3.** Permanova results for general differences in family composition of dispersed seeds by different ungulate species in Martinazo site and Bonferroni corrected pairwise comparisond between the four ungulate species ocurring in this site. Analysis has been conducted on seed content expresed both, as seeds per gram DW (column A) and as seed per fecal unit (column B). Significant differences in bold font

|  | Seed content |  |  |  |
| --- | --- | --- | --- | --- |
|  | A) Per unit DW (seeds.g<sup>-1</sup>) |  | B) Per fecal unit (seeds.deposition<sup>-1</sup>) |  |
| **Permanova** | Df=3, F= 2.142; **P=0.012** |  | DF= 3, F= 6.61; **P=0.0003** |  |
| **Pairwise comparisons** | F | P | F | P |
| Deer - Cow | **3.619** | **0.043** | **14.4** | **0.0006** |
| Deer - Horse | 2.137 | 0.345 | **9.88** | **0.0006** |
| Deer - Wild board | 0.774 | 0.998 | 1.534 | 0.9138 |
| Cow - Horse | 2.086 | 0.573 | 2.374 | 0.5976 |
| Cow - Wild board | 1.861 | 0.7596 | 3.531 | 0.1806 |
| Horse -Wild board | 1.441 | 0.457 | **2.852** | **0.0114** |

Similar results were obtained when NMDS analysis was conducted using data expresed per fecal unit, althought in this case a greather dispersion of points was observed along coordinates 1 and 2 (Figure 3b). As in previous analysis (Figure 3a), the *Plantaginaceae* family remained closely associatedwith cow. However, in this case several other families also showed strong associations with cow, including *Cyperaceae, Primulaceae, Poaceae, Cariophyllaceae* and *Polygonaceae*. Consistent with previous results, the *Ranunculaceae* family was again associated with deer, and additionally, the *Juncaceae* family showed a similar association in this analysis. *Fabaceae* and *Apiaceae* were primarily associated with horses, while no plant families were differentially associated with wild boar. PERMANOVA results (Table 3, column B) revealed significant overall differences in the taxonomic composition of dispersed seeds among ungulate species, with a higher level of significance than in the previous analysis (Df = 3, F = 6.61, P = 0.0003). Furthermore, a greater number of significant pairwise differences were detected, specifically between deer and cattle, deer and horse, and boar and horse.

### 3.3 Spatial patterns of seed dispersal

Spatial pattern of seeds dispersal (i.e., total number of seeds per fecal unit) produced different outcomes at the two sites under study (Figure 4). At Martinazo, a significant r-mark correlation function (P= 0.039, Figure 4a) indicates that seed content was influenced by the distance between fecal deposits. A positive effect of feces aggregation was observed at short distance (r = 0.5 to 2.5m), where the observed r-mark correlation function exceeded the upper envelop, while a negative effect was apparent at intermediate distances (r = 23.5 to 26.5 m), where it fell below the lower envelop. These results suggest a complex dependence of seed content on the spacing of fecal deposits. In Matasgordas, the observed r-mark correlation function fell within the simulation envelope across all scales, indicating no relationship between seeds content and the distance between fecal deposits (P = 0.340, Figure 4b). Schlater’s correlation function which assesses spatial covariance in seed content between two fecal deeposits separeted by a distance r, was also signifficant in Martinazo (P = 0.005, Figure 4c), exceeding the upper envelop at short distances (r = 0.5 to 2.5 m) and slightly above it at greater distances (r = 33.5 to 34.5 m). At Matasgordas, this function was non-significant (P = 0.595, Figure 4c), indicating no spatial dependence of seed content at any scale. The density correlation function which relates seed content in fecal deposits to nearby deposit density was marginally signifficant at Martinazo (P = 0.055, Figure 4e), exceeding the upper envelope at short (r = 0.5–2.5 m) and long (r = 47 m) distances. At Matasgordas, this function was non-significant (P = 0.285, Figure 4f).

**Figure 4.**
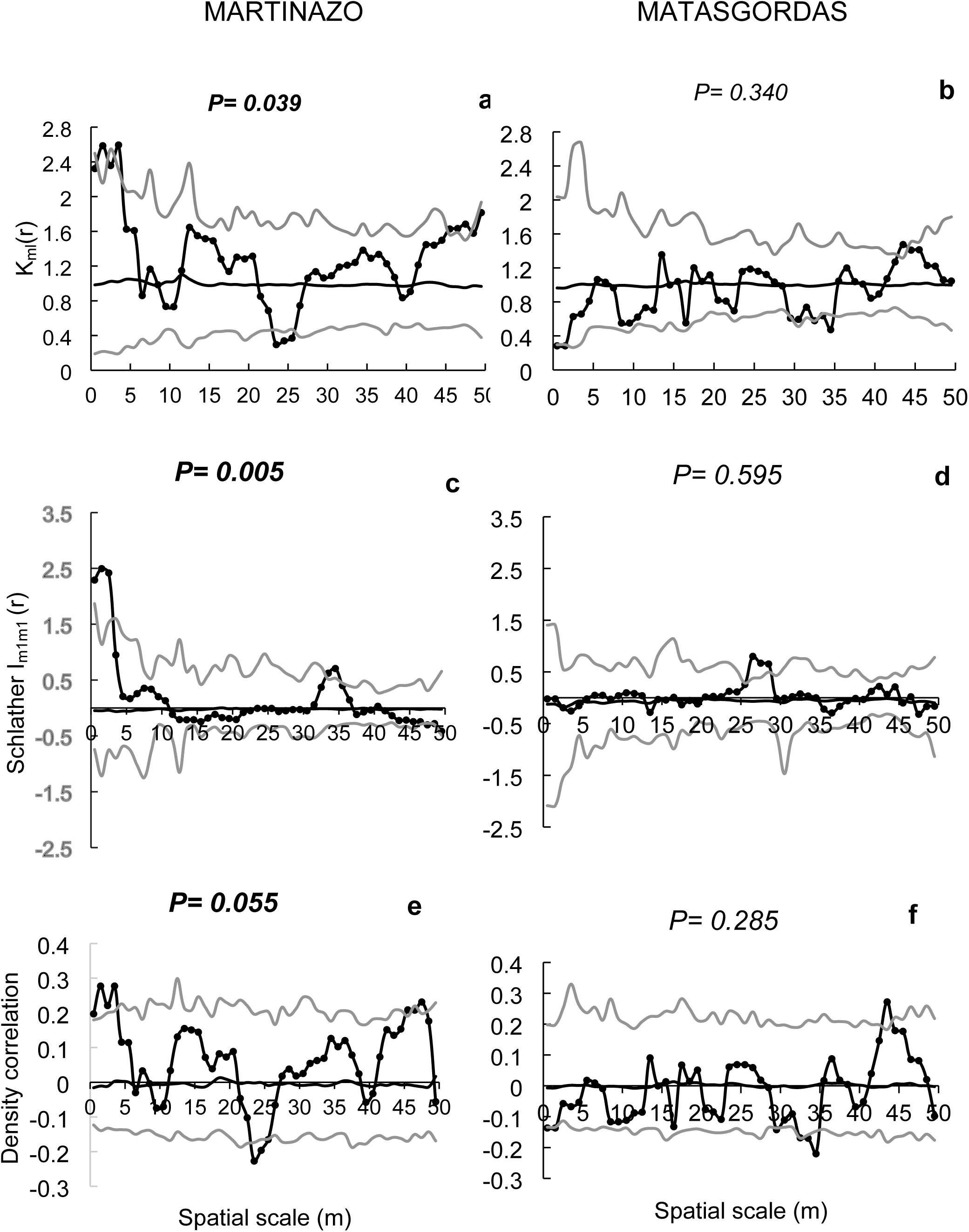
Mark correlation functions applied to to evaluate potential spatial structure in total number of dispersed seeds in Martasgordas (b,d,f) and Martinazo (a,c,e) sites. The r-mark correlation function (a, b) describes the mean number of seeds (mi) in a fecal unit at distance “r” of another unit. Schlather’s correlation function (c, d) quantifies the correlation between the number of seeds in two different units separated by distance “r”. Density correlation function (e, f) assesses the correlation between the number of seeds and the number of nearby feces located at a distance “r”. Observed functions= black line with solid dots. Expected functions under the null model of random mark i = black line. Corresponding simulation envelopes = grey lines (being the fifth lowest and highest values of the functions created by 199 simulations under random labelling). Goodness-of-fit (Baddeley et al, 2014) is used to test the overall fit of the random marking null model for the entire distance interval up to 50 m. Signifficant The *p*-value from GoF test (bold font shown in each panel) indicates departures of the observed function from the random marking null model over the distance interval of interest. Bold font = significant correlation funtion.

With regard to the plant families most frequently dispersed by ungulates at the two sites, we evaluated the potential spatial structure of seed dispersal for *Cyperaceae*, *Plantaginaceae* and *Cariophyllaceae* seeds in Martinazo (with occurrence frequencies of 44.7%, 36.3% and 36.0% respectively), and for *Valerianaceae*, *Fabaceae* and *Cyperaceae* seeds in Matasgordas (with occurrence frequencies of 48.3%, 38.1% and 37.3% respectively) (Figure 5 to 7).

**Figure 5.**
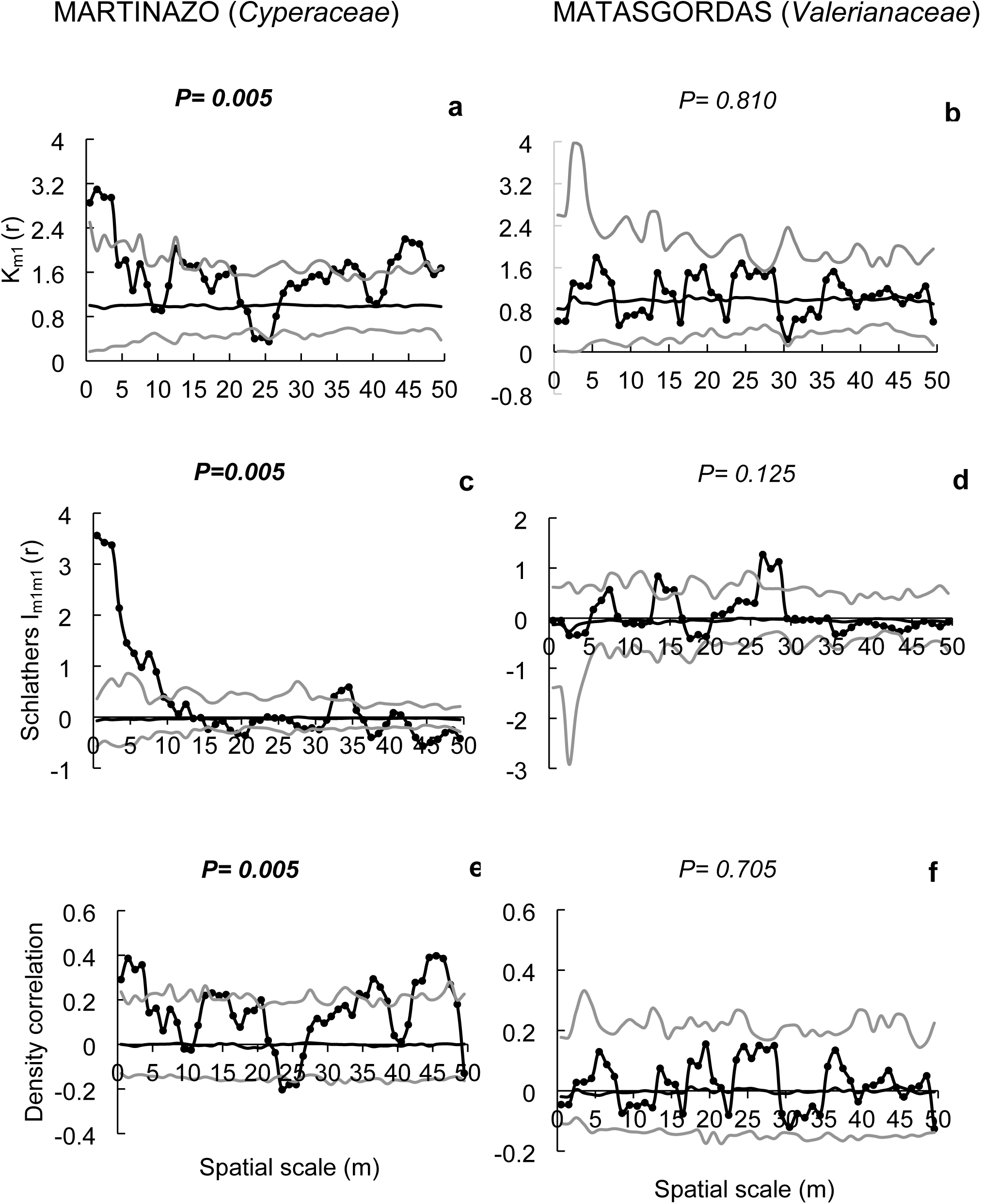
Mark correlation functions to evaluate potential spatial structure in the number of dispersed seeds of the most frequent plant families (indicated in bracket) in Martasgordas (b,d,f) and Martinazo (a,c,e). The r-mark correlation function (a,b) describes the mean number of seeds (mi) in a fecal unit at distance “r” of another fecal unit. Schlather’s correlation function (c, d) quantifies the correlation between the number of seeds in two different fecal units separated by distance “r”. Density correlation function (e, f) assesses the correlation between the number of seeds and the number of nearby feces located at a distance “r”. Meaning of the different coloured lines and symbols as in Figure 4.

The r-mark correlation function for the *Cyperaceae* seeds in Martinazo, was significant (P = 0.005, Figure 5a) indicating that the number of seeds per fecal unit was influenced by the distance between deposits. As in previous case (Figure 4a) there was a positive effect of feces aggregation on seed content (km (r) >1) at short distance (r = 0.5 to 3.5 m) where the observed function exceeded the upper envelop. Additionally, the function surpassed the upper envelope at longer distances (r = 42.5–46.5 m). The *Valerianaceae* seeds in Matasgordas, did not show a relationship between seed content and the distance between nearby feces (P= 0.81 Figure 5b). Schalater’s correlation function, which assesses spatial covariance in seed content between two fecal deposits separated by a distance r, was signifficant for *Cyperaceae* seeds in Martinazo (P= 0.005, Figure 5c). where the observed function greatly exceeded the upper envelope at short distance (r = 0.5 to 8.5m) and moderately at intermediate distance (r = 32.5 to 34.5 m). Conversely, this function was not signifficant for *Valerianaceae* in Matasgordas (P= 0.125; Figure 5d). Finally, density correlation function, which relates the number of dispersed seeds to the density of nearby feces was signifficant for *Cyperaceae* in Martinazo (P = 0.005; Figure 5e) exceding the upper envelop at short (r = 0.5 to 3.5 m) and long distances (r = 36.5 to 37.5 and 42.5 to 46.5 m), indicating a complex spatial dependence between seed content and nearby feces density. In contrast, this function was not significant for *Valerianaceae* in Matasgordas (P= 0.705; Figure 5f) with the observed function falling within the simulation envelope at all scales.

With respect to the *Plantaginaceae* family in Martinazo, a significant r-mark correlation function (P = 0.015, Figure 6a) indicates a spatial dependence between seed content and nearby feces distance. As in the previous case (i.e., *Cyperaceae* seeds; Figure 5a), a positive effect of feces aggregation (indicated by km(r) > 1) for *Plantaginaceae* in Martinazo was strongest at short distances (r = 1.5–3.5 m), where the observed function exceeded the simulation envelop. In contrast, the *Fabaceae* seeds in Matasgordas, did not show significant r-mark correlation function (P= 0.375; Figure 6b), indicating no relation between seed content and distance between nearby feces at any scale. The Schlather’s correlation function for *Plantaginaceae* seeds in Martinazo, also indicates significant covariance in seed content of nearby feces (P= 0.03, Figure 6c), and again the observed function greatly exceeds the envelop at short distance (r = 0.5 to 3.5 m). Conversely the non-signifficant Schlather’s correlation function for *Fabaceae* seeds in Matasgordas (P= 0.490; Figure 6d) indicates no relationship between number of dispersed seeds and the distance between feces. Finally, a signifficant density correlation function for *Plantaginaceae* seeds in Martinazo (P = 0.03; Figure 6e) suggests dependence between seeds content and nearby feces density. The density correlation function exceeded the upper simulation envelop at short distances (r = 1.5 to 3.5 m) and fell slightly below the lower envelop at intermediate distance (r = 23.5 to 25.5 m), reflecting again a complex spatial relationship. In contrast, the non-significant density correlation function for *Fabaceae* seeds in Matasgordas (P= 0.305; Figure 6f) indicates no dependence between seed content and nearby feces density.

**Figure 6.**
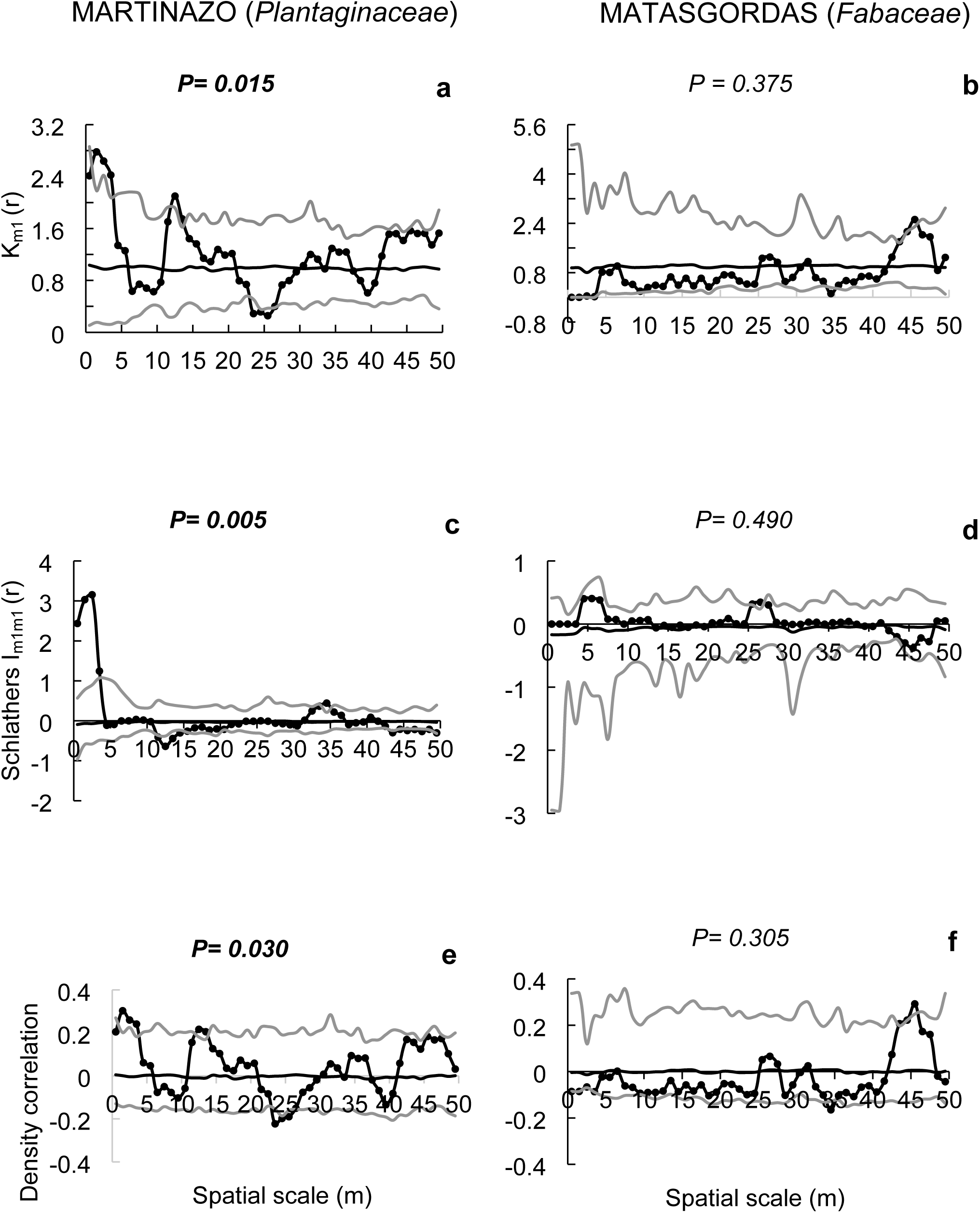
Mark correlation functions to evaluate potential spatial structure in the number of dispersed seeds of the most frequent plant families (indicated in bracket) in Martasgordas (b,d,f) and Martinazo (a,c,e). The r-mark correlation function (a,b) describes the mean number of seeds (mi) in a fecal unit at distance “r” of another fecal unit. Schlather’s correlation function (c, d) quantifies the correlation between the number of seeds in two different fecal units separated by distance “r”. Density correlation function (e, f) assesses the correlation between the number of seeds and the number of nearby feces located at a distance “r”. Meaning of the different coloured lines and symbols as in Figure 4.

Ultimately, non-significant r-mark correlation function for dispersed *Cariophyllaceae* seeds in Martinazo (P= 0.740, Figure 7a) and for *Cyperaceae* seeds in Matasgordas (P=0.80; Figure 7b) indicate that seed content in the fecal units was not influenced by the distance to nearby feces in either case. Similarly, non-significant Schalater’s correlation function for dispersed *Cariophyllaceae* in Martinazo (P= 0.385, Figure 7c) and for dispersed *Cyperaceae* seeds in Matasgorgas (P= 0.880, Figure 7d) show no spatial covariance in seeds content at any scale. Furthermore, density correlation function for *Caryophillaceae* in Martinazo and for *Cyperaceae* in Matasgordas were also not significant (P = 0.685; Figure 7e and P = 0.685; Figure 7f respectively) indicating that seed content in fecal units was not influenced by nearby feces density in either case.

**Figure 7.**
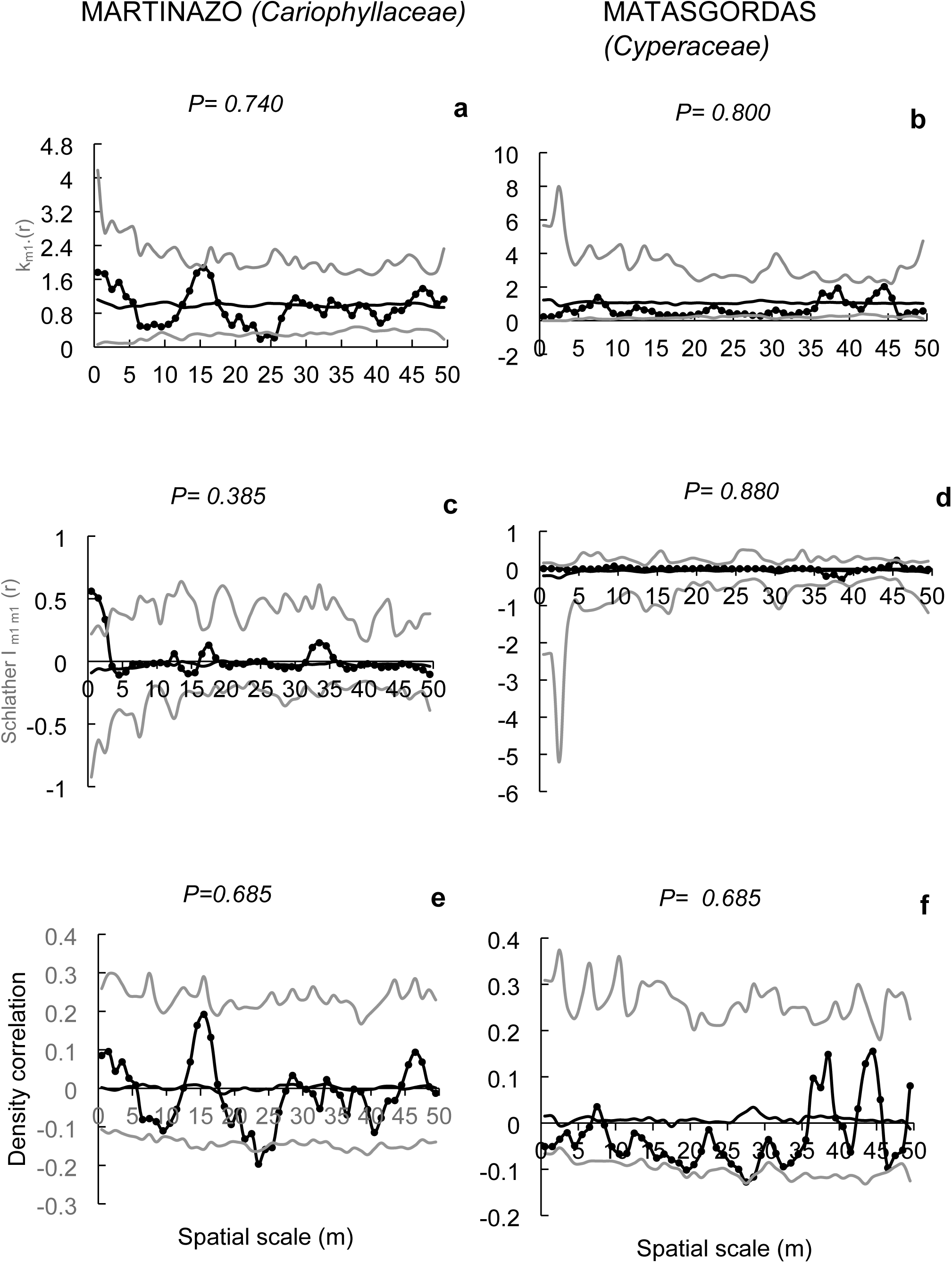
7 Mark correlation functions to evaluate potential spatial structure in the number of dispersed seeds of the most frequent plant families (indicated in bracket) in Martasgordas (b,d,f) and Martinazo (a,c,e). The r-mark correlation function (a,b) describes the mean number of seeds (mi) in a fecal unit at distance “r” of another fecal unit. Schlather’s correlation function (c, d) quantifies the correlation between the number of seeds in two different fecal units separated by distance “r”. Density correlation function (e, f) assesses the correlation between the number of seeds and the number of nearby feces located at a distance “r”. Meaning of the different coloured lines and symbols as in Figure 4.

## 4 DISCUSSION

### Effect of Site on Seed Dispersal

Previous studies report contrasting findings regarding the redundancy of seed species dispersed by sympatric ungulates. While some suggest significant overlap (Lawton & Brown, 1993; O’Connor & Crowe, 2005), others highlight clear interspecific differences (Polak et al., 2014; Picard et al., 2016). Our results reveal quantitative differences in herbaceous seed dispersal among grasslands in Doñana National Park. Seed deposition per gram of feces was more than twice as high at Matasgordas (12.4 seeds·g⁻¹) compared to Martinazo (5.7 seeds·g⁻¹; Table 2). However, when assessed per fecal unit, Martinazo showed higher seed loads (1,302 vs. 606 seeds·deposition⁻¹), likely due to differences in ungulate assemblages. Martinazo hosts large-bodied domestic ungulates (cattle and horses) with larger fecal deposits and higher seed loads (3,514 and 1,480 seeds·unit⁻¹, respectively), whereas Matasgordas is mainly occupied by deer, which disperse fewer seeds per deposition (300–615 seeds·unit⁻¹; Table 2). These findings underscore the influence of local ungulate composition on seed dispersal quantity (Bartuszevige & Endress, 2008).

In addition to quantity, seed taxonomic composition differed between sites. Dominant families at Matasgordas (*Valerianaceae*, *Fabaceae*, *Juncaceae*; Fig. 1a) were rare in Martinazo, where *Plantaginaceae*, *Cyperaceae*, and *Ranunculaceae* predominated. These differences likely reflect both variation in ungulate assemblages and inherent plant community heterogeneity, despite similar vegetation physiognomy of the seed arrival microsites (Table 1). Prior studies emphasize that seed content in feces is shaped by the local plant community (Malo & Suárez, 1995b; Picard et al., 2016). Though this study does not isolate ecological or anthropogenic drivers, NMDS and PERMANOVA analyses (Fig. 2) show significant taxonomic differences in deer-dispersed seeds between sites. Certain families (e.g., Valerianaceae, Fabaceae, Chenopodiaceae) were more common in Matasgordas, while others (e.g., Ranunculaceae, Boraginaceae) were more frequent in Martinazo. Such variation is expected in the heterogeneous “la vera” grasslands of Doñana (Lazo Contreras et al., 2001) and may be reinforced by local differences in ungulate density and composition (Gou et al., 2024).

### Ungulate species effect on seed dispersal

Our findings highlight the role of ungulate identity in shaping endozoochorous seed dispersal. Significant differences in dispersed seed families were observed among the four ungulate species at Martinazo (Table 3), independent of whether data were expressed per fecal unit or per gram of material. Cattle and deer, in particular, showed clear contrasts: *Plantaginaceae* was strongly associated with cattle, while *Ranunculaceae* was more frequently dispersed by deer (Fig. 3). Other associations included *Cyperaceae*, *Primulaceae*, and *Poaceae* with cattle, and *Polygonaceae* and *Boraginaceae* with deer, although these varied by the unit of measurement used (Figs. 3a, b). Additional differences emerged between deer, horse, and wild boar when data were expressed per fecal unit (Table 3B). These results align with prior studies showing low redundancy among sympatric ungulates (Bartuszevige & Endress, 2008).

Whereas most previous research links differences in seed dispersal to broad habitat preferences (Cosyns et al., 2005; Albert et al., 2015; Picard et al., 2016), our findings suggest that such differences can also manifest at fine spatial scales within a single habitat, the grassland under study. However, relatively small plot sizes and potential long-distance seed transport from adjacent habitats should be considered. Differences in digestive physiology and body size among herbivores likely contribute to dietary divergence and resulting seed composition (Couvreur et al., 2005; Traveset et al., 2007).

Selective ingestion also appears to play a role. Certain seed families, such as *Fabaceae* and *Caryophyllaceae*, occurred frequently despite low abundance, suggesting selectivity. This pattern was consistent across both sites. *Fabaceae* are known for their high nitrogen content, making them a preferred forage (Nisi et al., 2015; Siebert & Scogings, 2015; Han et al., 2020), and are frequently dispersed by ungulates and other herbivores, while *Caryophyllaceae*—along with *Fabaceae* and *Brassicaceae*—has been identified among the families most frequently dispersed by herbivorous mammals in Mediterranean woodlands (Malo & Suárez, 1995). Interestingly, more families exhibited this frequency-abundance mismatch in Matasgordas, indicating potential effects of ungulate density or other ecological variables. These findings emphasize the need for further research into plant selection mechanisms and their implications for dispersal processes.

### Spatial pattern of seed dispersal

Seed dispersal patterns differed markedly between sites. In Matasgordas, where deer were practically the only ungulates, no significant spatial structure was detected using Mark correlation functions (Fig. 4b, d, f), either for total seed content or for key families (*Valerianaceae*, *Fabaceae*, *Cyperaceae*; Figs. 5–7: b, d, f), indicating an absence of distance-dependent seed distribution. In contrast, Martinazo, with a more diverse ungulate community, exhibited clear spatial patterns. Short-distance aggregation of fecal deposits (0.5–2.5 m) was associated with increased total seed content and higher counts of *Plantaginaceae* and *Cyperaceae* (Figs. 4–6: a) while longer distances (23.5–26.5 m) showed negative associations. Schlather’s correlation function further revealed positive short- and long-range associations (33.5–34.5 m), while density correlation function, though marginally significant for total seed content (Figure 4e), showed strong significance for *Plantaginaceae* and *Cyperaceae* (Figures 5 and 6: e), indicating positive relationships at both short (0.5–2.5 m) and long (47 m) distances, with negative effects at intermediate distances (33.5–34.5 m). Collectively, these Mark correlation analyses consistently point to a positive short-range effect of fecal aggregation on total seed input and on *Cyperaceae* and *Plantaginaceae*, while suggesting more complex spatial interactions at broader scales.

These patterns likely stem from the presence of cattle, whose depositions are spatially clumped (Malo et al., 2000) and have the highest seed content among all the ungulates present (Table 2). Similar findings have been reported in other Mediterranean grasslands (Malo et al., 2000). Thus, high seed load combined with aggregated dung deposits likely drive the spatial structuring in seed dispersal observed in Martinazo. More complex patterns at broader scales may reflect environmental heterogeneity, such as temporary flooding or selective habitat use, which could influence fecal deposition and seed dispersal dynamics.

### Conclusions

Overall, our results highlight that seed dispersal by ungulates in Mediterranean grassland systems is neither random nor redundant. Instead, it is shaped by complex interactions between disperser identity, plant community composition, and landscape heterogeneity. These findings have important implications for biodiversity management in semi-natural and protected ecosystems and underscore the necessity of considering functional differences among herbivores when designing conservation and restoration strategies.

## ACKNOWLEDGMENTS

We thank the Doñana Research Coordination Office for field facilities. Dr Benito Valdés Castrillón and Dr Ádám Lovas-Kiss courteously collaborated in the identification of certain seed taxa. Marta Guerrero, Samuel Fuentes, Daniel Palma and Alejandro Rosales collaborated in sample processing.

## AUTHOR CONTRIBUTIONS

Conceptualization: María J. Leiva and José M. Fedriani

Formal analysis: María J. Leiva

Investigation and Methodology: María J. Leiva and José M Fedriani.

Resources and supervision: Jose M. Fedriani.

Writing – original draft: María J Leiva

Writing – review & editing: Jose M. Fedriani and María J. Leiva

